# cellPLATO: an unsupervised method for identifying cell behaviour in heterogeneous cell trajectory data

**DOI:** 10.1101/2023.10.28.564355

**Authors:** Michael J. Shannon, Shira E. Eisman, Alan R. Lowe, Tyler Sloan, Emily M. Mace

## Abstract

Advances in imaging, cell segmentation, and cell tracking now routinely produce microscopy datasets of a size and complexity comparable to transcriptomics or proteomics. New tools are required to process this ‘phenomics’ type data. Cell PLasticity Analysis TOol (cellPLATO) is a Python-based analysis software designed for measurement and classification of diverse cell behaviours based on clustering of parameters of cell morphology and motility. cellPLATO is used after segmentation and tracking of cells from live cell microscopy data. The tool extracts morphological and motility metrics from each cell per timepoint, before being using them to segregate cells into behavioural subtypes with dimensionality reduction. Resultant cell tracks have a ‘behavioural ID’ for each cell per timepoint corresponding to their changing behaviour over time in a sequence. Similarity analysis allows the grouping of behavioural sequences into discrete trajectories with assigned IDs. Trajectories and underlying behaviours generate a phenotypic fingerprint for each experimental condition, and representative cells are mathematically identified and graphically displayed for human understanding of each subtype. Here, we use cellPLATO to investigate the role of IL-15 in modulating NK cell migration on ICAM-1 or VCAM-1. We find 8 behavioural subsets of NK cells based on their shape and migration dynamics, and 4 trajectories of behaviour. Therefore, using cellPLATO we show that IL-15 increases plasticity between cell migration behaviours and that different integrin ligands induce different forms of NK cell migration.

## introduction

Imaging populations of cells over time with microscopy reveals behavioral heterogeneity and plasticity. Immune cells function in a variety of environments, and their morphology and motility is intrinsic to these functions. Many immune cell functions rely on cells tuning their morphology and motility behaviour, which allows them to probe, migrate, kill and remember (1–4). Advances in genetic and proteomics big data analysis techniques applied to biological data have enabled the stratification of subsets of cells using reduced dimensions. With the advent of fast, gentle microscopes and cell labeling dyes, and the development of AI-based software for segmentation and tracking, similar analyses can be applied to the morphological, motility and clustering features of cell images acquired using timelapse imaging. Here, we present a new Python-based tool, named cell plasticity analysis tool (cellPLATO), that separates heterogeneous populations of migrating cells into behavioural subsets and analyzes them through time. We apply cellPLATO to better understand the effect of IL-15 on natural killer (NK) cell migration on the integrin ligands intercellular adhesion molecule 1 (ICAM-1) and vascular cell adhesion molecule 1 (VCAM-1).

NK cells develop and function in tissues and blood, and their cell migration behaviours differ between circulating, adaptive, or tissue resident forms (5–7). Cell motility and morphology are key to the ability of NK cells to switch between developmental subsets and function as mature cells, in part influenced by integrin ligands in the microenvironment (6, 8, 9). NK cell maturation is characterized by an upregulation of β2 and β1 integrins (9), and immune cells in tissue secrete inflammatory cytokines upon infection which stimulate blood vessel endothelial cells to upregulate integrin ligands, especially ICAM-1 and VCAM-1. NK cells use lymphocyte function associated antigen 1 (LFA-1, αLβ2, CD11a/CD18) and very late antigen 4 (VLA-4, α4β1, CD49d/CD29) to bind ICAM-1 and VCAM-1 respectively, where integrin ligation induces shear flow resistant spreading, polarization and crawling migration along blood vessels which directly precedes transendothelial migration (10, 11). NK cells also use integrins to interact with the extracellular matrix (3) and integrin ligands expressed in tissues (12) and on other NK cells (13).

The expression level of ICAM-1 or VCAM-1 on endothelial cells or in tissues, or their cognate integrins LFA-1 and VLA-4 on lymphocytes, changes lymphocyte behaviour. In T cells, ICAM-1 expression induces migration against the direction of shear flow, whereas VCAM-1 induces migration in the direction of flow (14, 15). LFA-1 engagement with ICAM-1 on 2-dimensional glass surfaces induces widespread cell spreading and adhesion, commencing primarily at the cell’s front. In contrast, VLA-4 binding to VCAM-1 results in reduced spreading compared with ICAM1/LFA-1 and increased adhesion towards the rear of the cell. Both ligands activate partially overlapping signaling intermediates (16). LFA-1 forms nanoclusters that develop in the front of the cell then condense, becoming smaller in the middle of the cell and changing size with migration speed (12). While the effects on cell shape and migration speed of T cells migrating on ICAM-1 are distinct from those migrating on VCAM-1, whether ICAM-1 and VCAM-1 elicit different behaviours in subsets of mature NK cells is unknown.

The role of IL-15 in shaping how cells migrate on ICAM-1 vs VCAM-1 is not well elucidated. IL-15 is best known for its crucial role in the development, survival, activation, homeostasis, and killing capacity of NK cells (17–19), but IL-15 is also associated with integrin function and cell migration (20, 21). NK cells migrate more readily in response to IL-15, but it is unclear whether this response is due to chemotaxis to IL-15 or due to an indirect effect on cell surface receptors such as integrins (22). Stimulation of NK cells with IL-15 increases the expression of CD11a, the αL subunit of LFA-1 integrin, leading to increased cell adhesion to vascular endothelium (23). In T cells, IL-15 is associated with uropod formation, polarization, and increased adhesion to VCAM-1 or ICAM-1 (20). IL-15 expression is associated with chemokine expression (24) and its genetic deletion in mice is accompanied by downregulation of LFA-1 (25). Together, IL-15 affects cell migration through both genetic up-regulation and spatial reorganization of integrin molecules. Together, these result in cell shape change and influence cellular responses to ICAM-1 and VCAM-1.

Advanced microscopy, combined with new methods for image segmentation (26–29) and cell tracking (30–33), enables the analysis of thousands of cells over time, taking microscopy imaging into the realm of big data. We sought to develop a tool to measure and classify cell populations downstream of such methods. Existing tools for analyzing large populations of cells from microscopy data highlight the importance of morpho-kinetic analysis. Phenomic data can be used to distinguish macrophages from DCs and activated from non-activated neutrophils (34) by using tSNE for visualization and extracting example cells for visualization of characteristics (ACME) (35). Other methods address the need to link single cell visualizations and analysis to large datasets while analyzing diverse populations together (CAMI) (36) or use intracellular information to extract more parameters than segmentation alone (Pixie) (37). Others allow for the separation of cell subsets from heterogeneous populations (Traject3D (38) and CellPhe (39)) but require an intermediate level of coding knowledge to use.

Here, we present a new software tool (cellPLATO) that separates heterogeneous populations of cells into instantaneous behavioural subsets followed by behavioural trajectory signatures over time. The tool 1) makes measurements of morphology, migration and clustering per cell per timepoint, 2) uses dimensionality reduction (UMAP) and cluster analysis (HDBSCAN) to designate instantaneous behavioural clusters, 3) uses sequence similarity, UMAP and HDBSCAN to designate behavioural trajectories over time, and then 4) generates data visualization including de-abstractification of cells within each cluster and fingerprinting to compare between conditions. To apply this tool, we enriched NK cells from human blood, treated them with IL-15, and measured characteristics of cell migration and morphology over time on ICAM-1 or VCAM-1. cellPLATO analysis reveals distinct cell behaviour and response to IL-15 stimulation to uncover novel aspects of NK cell biology and highlight the utility of phenomics-based analysis for understanding the heterogeneity of cellular populations.

## Results

### Differential changes in morphology and motility of NK cells on ICAM-1 or VCAM-1

To study the migration characteristics of primary human NK cells, we isolated NK cells from peripheral blood using negative selection and confirmed by flow cytometry that >85% of cells were CD56+CD3/CD14/CD19 NK cells (Supp. Fig. 1). Cells were labeled with DNA dye and membrane dye for 30 minutes before plating onto an imaging chamber coated with ICAM-1 or VCAM-1 in the presence of IL-15 and imaging began after a 10-minute incubation (Fig. 1A). Cells were imaged by timelapse confocal microscopy to capture their migration on ICAM-1 or VCAM-1 (Fig. 1B and C). Following acquisition of imaging data from 2 healthy donors, NK cell masks were generated by segmentation with CellPose (27) and cells were tracked using btrack (31). Tracking and segmentation generated 14,825 and 6,584 distinct cell tracks for donor 1 and 2 respectively, with slightly more tracks identified in the VCAM-1 condition than the ICAM-1 condition (Fig. 1D). Cells were tracked for a minimum of 8 frames (320 seconds; 5.3 minutes) for an average of 164 frames (6560 seconds; 109.3 minutes) (Fig. 1E).

**Fig. 1.**
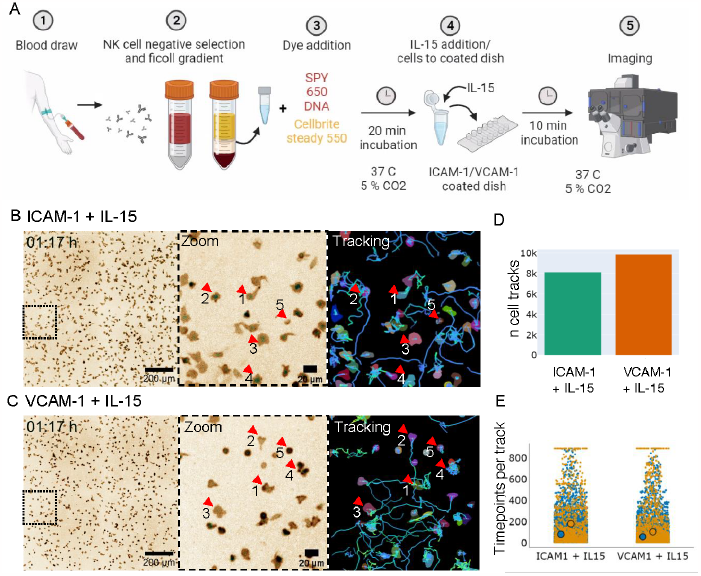
Morphological and migratory features of NK cells imaged, segmented, and tracked on ICAM-1 or VCAM-1 in the presence of IL-15. Freshly isolated human NK cells were pre-incubated briefly (10 min) with IL-15 then imaged on B) ICAM-1 or C) VCAM-1 surfaces in the presence of IL-15. Timelapse images of NK cells derived from PBMCs plated onto B) ICAM-1 or C) VCAM-1 coated glass slides. Dashed box (left) shows a zoomed region (middle) and example cell segmentation and tracking (right). Red arrows denote different morphologies of cells identified qualitatively as described in text. D) Number of cells detected per condition. E) Number of timepoints each cell was tracked. Blue = cells from donor 1, yellow: cells from donor 2. Large dots: mean per donor. n = 14,825 and 6,584 cells tracked respectively from two donors.

A diverse range of cell morphologies were identified qualitatively from the timelapse micrographs. Morphologically, NK cells on ICAM-1 exhibit polarized ‘teardrop’ shapes, semi-polarized shapes with wide lamellipodia, polarized shapes with multiple lamellipodia, or small and round shapes with no protruding edge (Fig. 1B, middle panel; red arrows). By extending this qualitative assessment to cell tracks, we noted that teardrop-shaped cells moved for long distances, while cells with wider morphology seemingly moved for shorter distances. Multi-lamellipodial cells were associated with frequent changes of direction, and small round cells were observed not migrating at all or beginning to migrate after long periods of quiescence (Fig. 1B, right panel; Movie 1 for raw timelapse; Movie 2 for segmentation and mask timelapse). When comparing micrographs of NK cells migrating on ICAM-1 with those on VCAM-1, we also observed behaviours that were shared between the two conditions but appeared at different frequencies. Morphologically, NK cells on VCAM-1 appeared less spread than on ICAM-1, but readily migrated in the presence of IL-15 (Fig. 1C). Timelapse movies indicate that some cells in the VCAM-1 condition were elevated at the uropod and lamellipodia, indicating adherence close to the back of the cell (Movie 1, Movie 2). The VCAM-1 condition generated small, rounded cells that appeared at a greater frequency than in the ICAM-1 condition (Fig. 1C).

Together, visual inspection of the micrographs, segmented masks, and cell tracks suggested that subpopulations of NK cells reacted differently to IL-15, ICAM-1, and VCAM-1. To further investigate cellular heterogeneity in our data, we developed a Python-based software tool designed to define sub-populations of cells by grouping their behaviours based on measurements of morphology and motility (described in detail in Methods). This tool (cellPLATO) calculates 29 measurements of morphology, motility, and clustering for each cell per timepoint. Certain measurements, such as cumulative length, are calculated over time windows as a series rather than instantaneously (between adjacent frames) or over all frames; this reduces measurement bias that can be introduced by cells entering or leaving the field of view. Here, a time window of duration 320 seconds (5 minutes 20 seconds) was used to optimize for the meaningful detection of cell behaviours while minimizing bias. We sought to understand basic population differences between NK cells migrating on ICAM-1 versus VCAM-1 and provide effective data visualization and statistics compatible with big microscopy data using cellPLATO.

### Rapid population-based changes in NK cell migration on ICAM-1 or VCAM-1 in the presence of IL-15

Having identified qualitative differences in cell morphologies between VCAM-1 and ICAM-1 conditions, we sought to understand how population-level metrics of cells were associated with single cell behaviors. cellPLATO outputs plots of difference (40) for each of the 29 morphological and kinetic metrics generated. Plots of difference are useful for large datasets where p-values become infinitesimally small due to sample size and do not necessarily reflect biologically relevant differences (41). We first investigated two fundamental measurements of cell migration and morphology, namely cell speed and cell area. When comparing conditions, the median migration speed of NK cells on VCAM-1 was 3.48 μm/min and 2.54 μm/min on ICAM-1 (Fig. 2A). The effect size distribution for VCAM-1 was greater, demonstrating statistical significance (p <0.00001) (42), and its distribution did not overlap with the control condition (ICAM-1). NK cells migrating on VCAM-1 also had smaller median cell area (114 μm^2^) compared with ICAM-1 (175 μm^2^) (Fig. 2B), with non-overlapping effect size distribution (p < 0.00001).

**Fig. 2.**
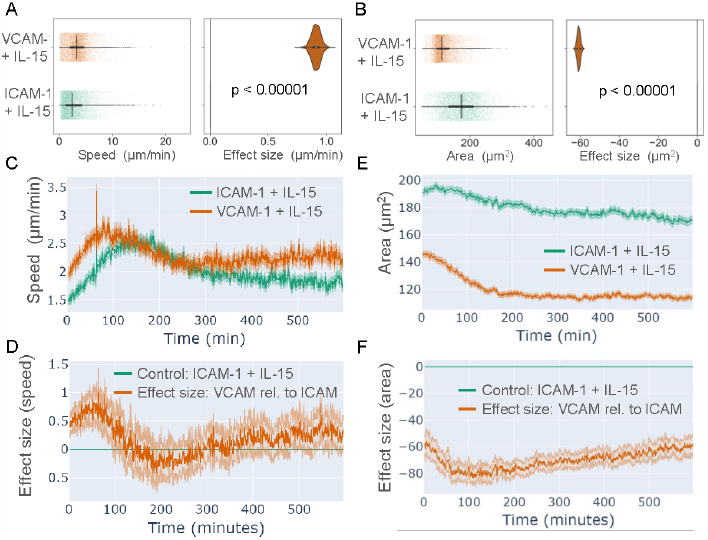
Features of NK cell speed and cell area on VCAM-1 and ICAM-1 in the presence of IL-15. Freshly isolated human NK cells were pre-incubated briefly with IL-15 then imaged on ICAM-1 or VCAM-1 surfaces in the presence of IL-15. Cells were segmented and tracked as in Fig. 1, and parameters of cell shape and cell migration were calculated using cellPLATO. A) Plot of differences (left) and effect size (right) for cell speed. B) Timeplot of mean cell speed over time, C) Timeplot of difference of effect size of cell speed of NK cells on VCAM-1 (orange line; VCAM-1 + IL-15) relative to control (blue line; ICAM-1 + IL-15). D) Plot of differences (left) and effect size (right) for cell area. E) Timeplot of mean cell area over time. F) Timeplot of difference of effect size of cell area of VCAM-1 (orange line; VCAM-1 + IL-15) relative to control (blue line; ICAM-1 + IL-15). n = 14,985 cells from 1 donor.

To define the acute effect of IL-15 on NK cells migrating on ICAM-1 or VCAM-1, the mean speed per timepoint was calculated and plotted (Fig. 2C). Mean speeds of cells migrating on ICAM-1 and VCAM-1 both increased in response to IL-15 addition. Cells on VCAM-1 reached a peak mean migration speed of 2.82 μm/min at 82 minutes before decreasing to consistently 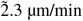 at 290 minutes until the end of imaging at 600 minutes. Cells on ICAM-1 took longer to reach a slightly lower peak mean migration speed of 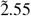 μm/min at 120 minutes before decreasing to a plateau of <2 μm/min at 310 minutes until 600 minutes. Visualization by plots of difference over time allowed us to statistically measure the difference over time between ICAM-1 as a reference condition and VCAM-1 by calculating and plotting the mean effect size at every time point. NK cells on VCAM-1 plated were significantly faster than ICAM-1 cells for the first 100 minutes reaching a peak mean difference of 1.02 μm/min, after which they were not statistically dissimilar (Fig. 2D).

Cell area was found to be consistently higher over time in NK cells migrating on ICAM-1 than cells on VCAM-1. NK cells on ICAM-1 had a continuous decrease in mean cell area beginning at 193 μm^2^ and ending at 174 μm^2^ (Fig. 2E). NK cells on VCAM-1 had a rapid decrease in area in the first 150 minutes, beginning the experiment with a mean area of 142 μm^2^ and reaching a mean area of <120 μm^2^ at 150 minutes, which remained consistent throughout the remainder of the experiment. Plots of difference over time revealed a consistent statistically significant difference in cell area of -60 μm^2^ in VCAM-1 relative to ICAM-1 cells throughout the experiment (Fig 2F). Cells were still migrating at the end of the experiment, indicating that changes we observed were in response to exposure to ICAM-1 or VCAM-1 and were not due to phototoxicity during imaging.

In summary, IL-15 treatment led to increased speeds of NK cells migrating on VCAM-1 or ICAM-1 and cells reached maximum speeds more rapidly on VCAM-1. Cells on ICAM-1 had consistently larger cell areas than cells on VCAM-1, which experienced a significant decrease in cell area in the first 150 minutes of the experiment. Together these data demonstrate that NK cells have different features of migration in response to IL-15 that is dependent on integrin ligands.

### NK cells migrating on ICAM-1 or VCAM-1 can be grouped into eight distinct ‘per-timepoint’ behavioural clusters using UMAP and HDBSCAN

While population-based averages identified differences in cell speed and area, they did not stratify subpopulations of cells responding differently to IL-15, ICAM-1, or VCAM-1. Additionally, single metrics cannot capture the complex nature of single cell behaviour, so we sought to group cells based on multiple metrics measured between adjacent timepoints. cellPLATO performs UMAP on morphological/motility parameters then uses HDBSCAN cluster analysis to define behavioural clusters. Exemplar cells that best represent each cluster are displayed as single cell graphics to de-abstractify the behaviour that each cluster represents, and the top three contributing metrics are displayed for each behaviour type. Finally, the median values and main characteristics for the most contributory metrics for each cluster are displayed in a table.

We performed dimensionality reduction and cluster analysis on our combined dataset of NK cells migrating on VCAM-1 or ICAM-1 after IL-15 activation, and plotted UMAPs 1, 2 and 3. Each point represents a single cell’s behavioural status at each time point, and cells belonging to each cluster identified using HDBSCAN are differentiated by colour (Fig. 3A). 8 separate behaviours (clusters) of cells were identified over the range of UMAP and HDBSCAN hyperparameters tested. To explore the quality of clustering, we calculated the silhouette score, a measure of how well a given datapoint has been classified in a cluster (see Methods), which we computed as 0.84 (range: -1 to 1). CellPLATO gives users the option to output several cluster scoring metrics, but such scoring methods are highly dependent on the nature of the data. We therefore combined scoring the clusters with visualization of cell contours from each cluster and the top 3 contributory metrics to their cluster membership (Supp Fig. 2). Cells belonging to the same cluster appeared proximal in dimensionally reduced space, were highly visually similar, and shared similar morphological and migratory characteristics. We also visualized the contribution of each metric as a heat map in UMAP space (Supp. Fig. 3). Metrics contributed to different extents to different clusters, indicating that they were useful in separating cells into behavioural subsets.

**Fig. 3.**
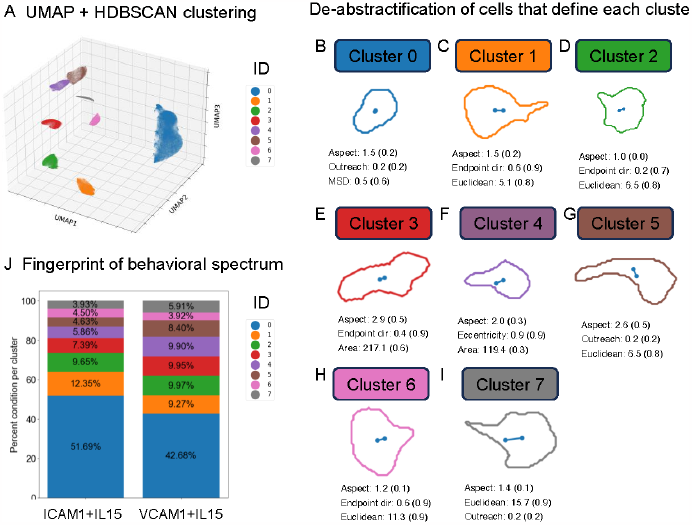
8 distinct behaviours disambiguated from IL-15 treated NK cells migrating on ICAM-1 or VCAM-1. A) UMAPs 1, 2, and 3 followed by HDBSCAN cluster analysis identified 8 clusters of distinct cell behaviours based on morphological and motility-based metrics. B-I) De-abstractification of the identity of cells in each cluster with exemplar cell graphics representative of each cluster accompanied by the metrics that best defined their membership to that cluster. Absolute values for each metric are followed by scaled values in parentheses. J) Representation of each cluster ID by condition. n = 14,985 cells from 1 donor.

To intuitively display cell migration and morphology behaviours, cellPLATO graphically displays exemplar cells from each cluster beside their 3 most contributory metrics that define cluster membership (Fig. 3B to I). Cells in clusters adopted a range of behaviours driven by distinctive combinations of aspect ratio, endpoint directionality, outreach ratio, MSD, eccentricity, and area. To understand the distribution of these de-abstractified per timepoint cell migration behaviours on VCAM-1 or ICAM-1, the percentage of cells found in each behavioural cluster was calculated to produce a behavioural ‘fingerprint’ for each condition (Fig. 3J). Using this analysis, clear proportional differences in the spread of the 8 behaviours emerged when comparing NK cells interacting with VCAM-1 compared with ICAM-1 (Fig. 3J), with NK cells from ICAM-1 or VCAM-1 conditions occupying all 8 of the clusters in different proportions (Supp. Fig. 4). To complement the graphics and the fingerprint plot, the defining characteristics for each cluster are described in Table 1, and median values are available in Supplementary Table 1.

**Fig. 4.**
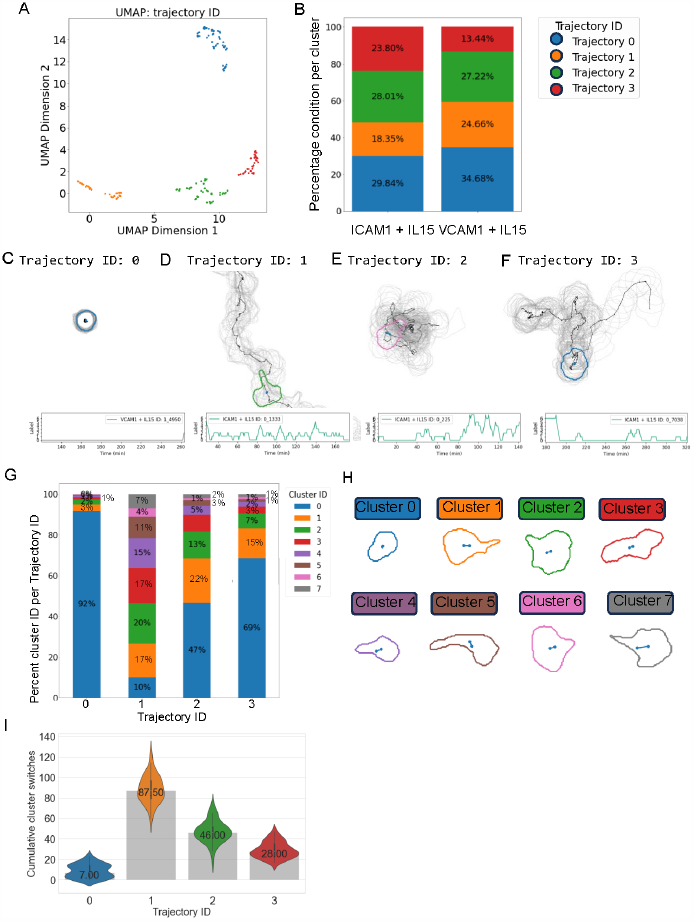
Behavioural trajectory cluster analysis reveals differential trajectories of NK cells on ICAM-1 or VCAM-1. Trajectory analysis was generated by pairwise similarity analysis of time sequences of cluster IDs. A) UMAP plot coloured by trajectory ID. B) Fingerprint plot showing the frequency of trajectory ID by condition. C-F) Graphical representations (top) and time plots of behavioural cluster ID (bottom) of each trajectory ID. G) Fingerprint plot showing the frequency of single timepoint behavioural cluster IDs for each trajectory ID. H) Graphical depictions of single timepoint behaviours represented in G (also shown in Fig. 3). I) The plasticity of cells in each behavioural trajectory ID measured by the median number of cluster switches. J) Mean number of cluster switches over time. n = 187 cells filtered to include only cells with 200 to 220 timepoints randomly selected from 14895 cells from 1 donor

Comparing the ICAM-1 and VCAM-1 conditions, most cells in both conditions were associated with behavioural cluster 0 and were cells with elliptical morphology that moved back and forth in place (Fig. 3J: percentages, Fig. 3B: graphical representation). The most contributory factors to cluster 0 cells were a medium aspect ratio (1.5), a medium outreach ratio (0.2) and a low MSD (0.5 AU) (median values are shown in Supp. Table 1). Cells with behaviours 0 and 1 were more prevalent in NK cells on ICAM-1 compared with VCAM-1. Cluster 1 cells were semi-polarized cells that underwent a directed move to a new location (Table 1, Fig. 3C). A similar proportion of cells on ICAM-1 and VCAM-1 adopted semi-polarized morphology with high migration, denoted by behavioural cluster 2 (Fig. 3D). Clusters 3, 4 and 5 were increased in proportion in VCAM-1 compared to ICAM-1 and constituted highly polarized cells with medium to high migratory displacements (Fig. 3E-G). Cluster 6 cells were increased in ICAM-1 versus VCAM-1, and represent cells that have high polarization, straightness and Euclidean distance and are thus highly migratory (Fig. 3H). Cluster 7 cells were more prevalent on VCAM-1 and represented highly polarized cells that made large displacements, but which had low outreach (Fig. 3I). A cell with high Euclidean distance but low outreach denotes those that have irregular patterns, stops or circular/cyclic paths that result in a limited maximum distance relative to Euclidean distance, ending close to their start point. In summary, cells on VCAM-1 had different frequencies of behavioural cluster residency when compared with ICAM-1. ICAM-1 induced the prevalence of NK cells defined by medium polarization, medium displacement and directed medium to fast migration, whereas VCAM-1 induced more extreme polarization and faster movement, with cells more frequently returning close to their initial position.

### Behavioural sequence analysis reveals stationary, confined, oscillatory and exploratory sub-populations of cell behavioural trajectories through time

To see how behaviours of cells at single timepoints correlated with their behaviour over time, we performed behavioural trajectory analysis. Every cell has a behavioural signature at each timepoint defined by a cluster ID. Therefore, single cell tracks can also be converted to a number sequence reflecting changes in behaviour over time as they switch between cluster IDs. CellPLATO implements sequence similarity analysis using Damerau-Levenshtein (normalized edit distance) (43) to perform pairwise comparison of trajectories of all cell tracks as they change over time. Similarity scores are used with UMAP and HDBSCAN to group sequences of similar sequences and to designate each type with a ‘trajectory ID’. While normalization helps reduce the influence of sequence length, sequences of vastly different length cannot be meaningfully compared by edit distance metrics. Therefore, analysis was performed on a filtered subset of 187 cells with similar track lengths in time (200-220 timepoints). We confirmed that this dataset maintained the same behavioural trends between conditions as the full dataset (Supp. Fig. 5). For clarity, we refer to trajectory IDs to denote cells that have similar strings of behaviours over time and refer to single timepoint cluster IDs to refer to the behavioural clusters identified previously.

**Fig. 5.**
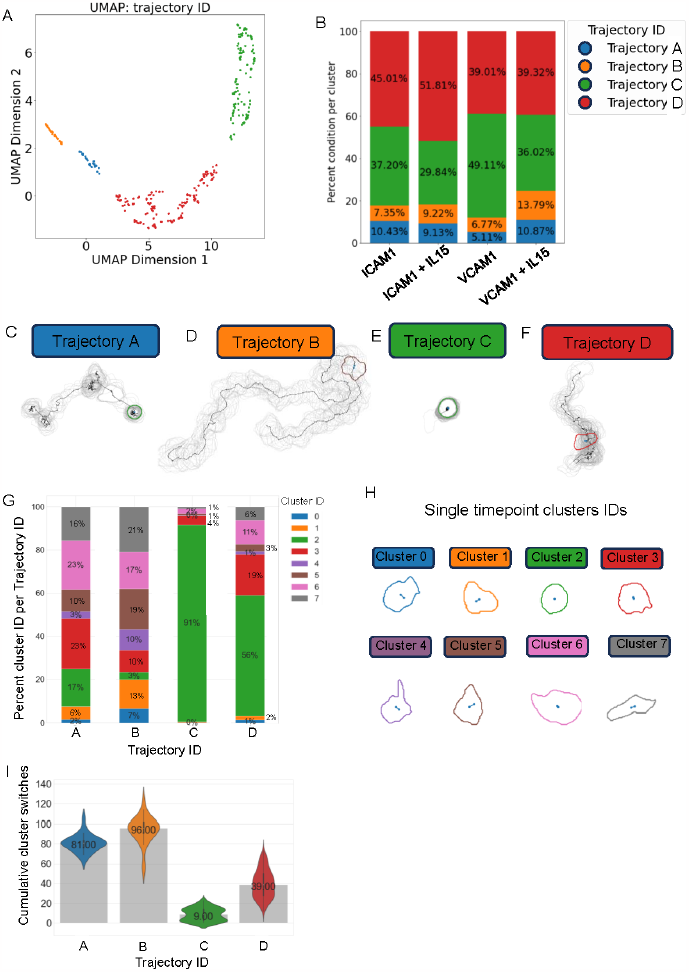
Effect of IL-15 on behaviourally distinct NK cell subsets migrating on ICAM-1 or VCAM-1. A) UMAP plot coloured by trajectory ID. B) Fingerprint plot showing the percent spread of each trajectory ID per condition. C to F) Graphical representations (top) and the behavioural cluster ID over time (bottom) for trajectories 0 to 3. G) Fingerprint plot showing the percent spread of single timepoint behavioural cluster IDs for each trajectory ID. H) Graphical depictions of single timepoint behaviours represented in G. I) The plasticity of cells in each behavioural trajectory ID, measured by the median number of cluster switches. J) The mean number of cluster switches over time. n=187 cells, filtered to include only cells with 200 to 220 timepoints randomly from 22,370 cells from 1 donor.

Sequence similarity analysis identified 4 distinct behavioural trajectories with clear separation in UMAP space (Fig. 4A). Measuring the distribution of trajectories per condition showed that trajectories were found in different frequencies when comparing NK cells on ICAM-1 vs VCAM-1 (Fig. 4B). Specifically, the proportion of trajectory 0 and trajectory 1 cells was reduced on ICAM-1 compared with VCAM-1 (Fig. 4B; blue and orange bars respectively) while trajectories 2 and 3 were increased (Fig. 4B; green and red bars respectively).

To understand the behavioural nature of each trajectory type, we de-abstractified the trajectory ID clusters by pulling exemplar trajectories from UMAP space. These were displayed as graphics denoting their contour and track through time with their single timepoint cluster ID over time (Fig. 4C to F). Trajectory 0 cells appeared visually stationary and morphologically round from visual inspection of the contour (Movie 3). The time-plot showed these cells occupied mostly single timepoint cluster 0, representing a stationary trajectory type (Fig. 4C). Trajectory 1 cells appeared very migratory and had directed migration with frequent switching between migratory behaviours and cell morphologies (single timepoint clusters 1 to 7), representing an oscillatory/exploratory trajectory type (Fig. 4D; Movie 4). Trajectory 2 cells transitioned from a stationary, rounded phenotype before switching to a highly migratory mode in a confined area (Fig. 4E; Movie 5). Trajectory 3 cells also transitioned between stationary rounded or slightly eccentric exploratory phenotypes to highly migratory ones, but did so in short bursts, moving to wider areas in short oscillatory bursts (Fig. 4F; Movie 6). To further validate verify the graphical exemplars shown, additional examples of each trajectory (Supp. Fig 6) and their cluster ID over time (Supp. Fig 7) were plotted and their similarity was confirmed with a silhouette score of 0.76.

To quantify the behaviours making up cells belonging to each trajectory ID, the composition of each trajectory ID by single timepoint cluster ID was plotted (Fig. 4G). The single timepoint exemplar cell graphics representative of each of the single timepoint cluster IDs from Fig. 3 are shown again in Fig. 4H and can also be found Table 1. Together, the plot and graphics show that each trajectory ID is made up of different proportions of identified single timepoint behaviours (Fig. 4G, H). As suggested by the qualitative observations described above, trajectory 0 cells primarily occupied cluster ID 0, remaining circular with low migration (Table 1). Trajectory 1 cells spent time in all single timepoint cluster IDs, but especially those associated with medium speed and varying levels of cell polarization and spread indicating a high level of shape change and exploratory migration. Trajectory 2 cells were characterized by a 47% membership in cluster ID 0, indicating roundness with low migration, coupled with 22% and 13% membership in moderately migratory clusters 1 and 2, and the rest of the time spent in highly migratory clusters 3, 4, 5, 6 and 7. As shown by timeplots (Fig. 4E), trajectory 2 cells appeared to move through clusters sequentially, from a state of low migration to a state of higher migration later in time, suggesting a delayed activation followed by high activity. Trajectory 3 cells spent 69% of their time as rounded, low migration cluster 0 cells, and had <15% membership in all other clusters, indicating short oscillatory bursts of migratory activity (Fig. 4G).

The fingerprint of single timepoint behaviours that make up each trajectory defined the cluster ID composition of each trajectory ID (Fig. 4G) but did not describe whether cells remained in each cluster ID for long periods of time or switched between clusters quickly. To quantify these changes, we calculated the number of single timepoint cluster ID switches made by each cell (Fig. 4I). Trajectory 0 cells make a median of 7 switches over their time course (200 to 220 frames; 133 to 146 minutes), trajectory 1 cells make 87.5 switches, trajectory 2 cells make 46, and trajectory 3 cells make 26. Together this indicates that the trajectories we identified have distinct behavioural plasticity.

In summary, trajectory analysis revealed 4 subpopulations of cells identified by clustering similar strings of single time-point behaviours (cluster IDs) in feature space, namely a stationary subset (trajectory 0), and highly plastic exploratory subset (trajectory 1), and delayed activation subset (trajectory 2), and an oscillatory confined migration subset (trajectory 3). All trajectories were represented on ICAM-1 and VCAM-1 surfaces. However, on ICAM-1, the oscillatory confined migration subtype was much more prevalent than on VCAM-1, where the highly plastic and exploratory subset was more prevalent.

### IL-15 acutely changes the migration and morphology of NK cells on integrin ligands

Given the highly migratory behaviour of NK cells migrating on ICAM-1 or VCAM-1 in the presence of IL-15, we wanted to determine the contribution of IL-15 to cell migration behavior relative to integrin ligands alone. We imaged NK cells migrating on ICAM-1 or VCAM-1 in the presence or absence of IL-15 in four wells simultaneously using multipoint montage images at 20X magnification for 600 minutes with 40 seconds between time-points. We then carried out a full cellPLATO analysis, including 1) measurement of cell characteristics per timepoint, 2) UMAP and HDBSCAN clusters of single timepoint behaviours (assignment of cluster IDs), 3) de-abstractification of those behaviours, 4) sequence similarity analysis, UMAP and HDBSCAN to produce trajectory IDs and 5) analysis of the plasticity of those trajectory IDs. This new dataset consisted of 22,370 cell tracks derived from a third healthy donor. It is important to note that while the same set of 8 single timepoint behaviour types were detected, and the same set of 4 trajectory types were extracted, they are displayed in a different order as this is a completely new analysis that does not share the same feature space as previous data. However, identification of the same set of trajectory clusters in both analyses is a good indicator that the tool works reliably to stratify the same behaviorally distinct subsets of cells from heterogeneous data and that these patterns of migratory behavior were conserved between NK cells from healthy individuals. We label the trajectories A, B, C and D to differentiate them from those identified in the previous analysis.

For this analysis, we moved directly to designating trajectory IDs generated by comparing the similarity of sequences of behaviours identified as single timepoint cluster IDs. Data were first filtered to include only track lengths of 200 to 220 timepoints before plotting their UMAP coordinates colored by trajectory ID, showing 4 well separated clusters (Fig. 5A). We then plotted the percentage occupancy of each trajectory type per condition (Fig. 5B), followed by graphically representative examples of each trajectory ID (Fig. 5C-F). Graphics showing the nature of the trajectories are intuitive and allowed us to quickly match phenotypes with previously identified trajectories. Here, trajectory A cells are oscillatory confined cells, trajectory B represents exploratory and very plastic cells, trajectory C represents stationary cells, and trajectory D represents cells with delayed activation. Comparing the distribution of trajectories between conditions, trajectory A cells (Fig. 5B, blue bars) make up a similar proportion of the population when comparing NK cells treated with IL-15 compared with those untreated with IL-15 on ICAM-1. On VCAM-1, the proportion of trajectory A cells doubles following treatment with IL-15 (Fig. 5B). Trajectory A cells are frequent shape changers and tend to move small distances, with frequent arrests with long dwell times (grey contours and black track line; Fig. 5C). Trajectory B cells increase slightly their proportion in the presence of IL15 on ICAM1 and their frequency is also doubled in the presence of IL-15 on VCAM-1 (Fig. 5B, orange bars). These cells migrate long distances, making very few stops, and have highly spread morphology (Fig. 5D). Trajectory C cells are decreased in NK cells on ICAM-1 and VCAM-1 when IL-15 is present (Fig 5B, green bars). There is a much higher proportion of trajectory C cells on VCAM-1 compared with ICAM-1. Such cells represent very round and stationary cells with smaller morphological changes (Fig. 5E). Trajectory D cells are increased in the presence of IL-15 on ICAM-1 but retain the same proportion in the absence and presence of IL-15 on VCAM-1 (Fig. 5B, red bars). Trajectory D cells have long periods of time round and stationary before experiencing bursts of activity with sustained migration and fast morphological change (Fig. 5F).

We further quantitatively calculated the percentage proportion of each single timepoint cluster ID for each of the 4 trajectories (Fig. 5G). This was combined with graphical exemplar representations of the single timepoint cluster IDs (Fig. 5H). Highly plastic trajectory A cells occupied all 7 single timepoint clusters at a different proportion to trajectory B cells, indicating that these small changes differentiate the two subsets. Most noticeably, trajectory B cells occupy cluster 5 and 7 (Fig. 5H, brown and grey contours) to a much higher proportion: two highly spread phenotypes with highly directional migration; the time spent in these cluster types appears to drive their considerably longer tracks and greater proportion of time spent with a large footprint. Trajectory C cells mostly occupy single timepoint cluster 2 (rounded, stationary cells), though importantly their switch to clusters 3 and 6 indicates that they are not dead, they are just not migratory even in the presence of IL-15. Trajectory D cells also spend long periods of time in a stationary rounded state (cluster 2) but are different to trajectory 2 cells because they then transition into migratory behaviours later in their track.

We further compared the different trajectories by plotting how they were driven by switches each cell makes between clusters over the time of imaging used to generate trajectories (200 to 220 timepoints) (Fig. 5I). Trajectory A cells and trajectory B cells were the most behaviourally plastic subsets of cells, making median 81 and 96 behavioural switches over the timecourse (median). This was starkly different to trajectory C cells, which made only 9 switches, and trajectory D cells, which made an intermediate 39 switches.

Together, these analyses show that a switch to more migratory behaviour happens in direct response to IL-15 addition. Using a data-driven approach that allows us to compare the spread of behaviourally distinct subsets of cells in a heterogeneous population, we show that the increase in migratory behaviour happens in a specific subset of the cells. Further, we confirm that 4 different modes of migratory and morphological behaviour are present in purified NK cells isolated from human blood. These include a stationary format unresponsive to IL-15, a format with delayed activation, a format that is highly plastic and exploratory, and a format that oscillates between short bursts of migration and arrest. Finally, we show that VCAM-1 favours the highly plastic form of migration, where ICAM-1 favours short bursts of oscillatory migration or delayed activation followed by steady migration.

## Discussion

Advanced imaging methods, cell segmentation (26–29, 44) and cell tracking (30–33) technologies now allow us to capture behavioural heterogeneity of large populations of cells from microscopy data. However, new analysis approaches are required to extract human-understandable biological concepts from such large and complex data. Traditional analysis tools typically reduce rich datasets to population-level averages where tracking and morphology metrics are averaged across different conditions. Such analyses effectively average out the heterogeneity, ignoring the contribution of different subpopulations of cells. Here we introduce cell plasticity analysis tool (cellPLATO), a simple Jupyter notebook-based platform designed to embrace the heterogeneity present in biological microscopy data.

CellPLATO proceeds in several steps, namely 1) measurement, 2) UMAP/HDBSCAN timepoint behaviour cluster designation, 3) behavioural trajectory cluster designation and 4) full visualization including de-abstractification cells within each cluster and fingerprinting to compare conditions. Briefly, the tool takes as input csv or h5 files containing tracking and segmentation data from btrack (31), TrackMate (45) or Usiigaci (28). Data should be organized in a two-tiered hierarchy, with separate folders for conditions at the top level, and subfolders for experimental replicates within each condition. After collation of data from multiple conditions and replicates, cellPLATO calculates 29 migration and morphology metrics per timepoint per cell, generating plots of difference and plots of difference over time automatically for population averages of all metrics per condition and saving them in a folder. CellPLATO then employs UMAP and HDB-SCAN (based on UMAP or raw data) to separate the cell population into subpopulations based on their morphological and motility-based features per timepoint, giving each cell at each timepoint a cluster ID. To explain the tangible nature of these clusters, HDBSCAN identifies exemplars which cellPLATO plots as graphics representing individual cells. The metrics that most contribute to the membership of each cluster based on their maximal variation from other clusters is listed next to each exemplar. To validate the data, clustering scores are extracted, and many exemplar cells from each cluster are visualized in a grid format. Having validated that single time-point behaviour ID designations for every cell at every time-point are reliable, the list over time provides a behavioural signature. CellPLATO then uses Damerau-Levenshtein distance followed by a second layer of UMAP/HDBSCAN to group together similar behavioural sequences. Using this method, cellPLATO can identify subpopulations of changing cell behaviours over time or behavioural trajectories. Trajectory cluster exemplars are extracted from the population data and displayed graphically, including as movies showing cluster change over time. These are coupled to static outputs to show the whole cell track and segmentation, as well as timeplots of cluster membership. cellPLATO also generates a ‘fingerprint’ plot, representing percentage trajectory cluster distributions between conditions and defining the single timepoint cluster ID makeup of each trajectory ID. Coupled with graphical understanding of the nature of each trajectory, experimenters can intuitively understand which cell behaviors are the most and least prevalent in each condition. Finally, cellPLATO computes the cumulative plasticity of cells belonging to each trajectory ID, a measure of the number of single timepoint cluster switches performed by each cell over time.

Taken together, CellPLATO offers a platform for comprehensively analyzing heterogeneous segmentation and tracking data. The use of a single config file for input parameters such as pixel size simplifies analysis for those with limited coding experience, and the workflow unfolds automatically, either with a single command line or via modular Jupyter notebooks. The tool is optimized for handling large datasets, taking 3 hours of computing time on a mid-range CPU to analyze the 40,568 cell tracks in this manuscript, coupled with all visualization. The codebase itself is highly modular and adaptable should users wish to add new metrics prior to downstream analysis.

### Comparison to existing tools

Recently, multiple tools including Traject3D (matlab) (38) and CellPhe (R) (39) have been developed to identify subsets of cells within complicated microscopy data. These tools can be challenging to use for individuals with less coding expertise, are coupled to specific tracking file formats, and do not provide intuitive data visualization for linking population level heterogeneity to biological understanding of single cell behaviour. cellPLATO improves on each of these aspects, as users can use one of three tracking algorithms to input their data by simply noting the input format and master folder in a config file. The three input formats are highly generalizable, and so can be used for other tracking algorithms outside of those mentioned. A jupyter notebook allows a modular approach to the analysis, providing the user with clear prompts and giving explanations as to what is happening at each step. Alternatively, the workflow can be run through in its entirety using a single command line. Using dimensionality reduction and cluster analysis quickly abstractifies data, however cellPLATO combats this by providing mathematically representative single cell analysis coupled to the population analysis in the form of fingerprint plots. CellPLATO also improves upon existing tools by implementing graphical visualization of statistics in the form of effect size plots. Existing tools that work on phenomics data often use traditional p-value based statistical testing only, often producing tiny P-values for insignificant biological changes (41, 46). To address this cellPLATO computes the effect size distribution between conditions, providing a clearer perspective on differences compared to a control condition (based on an R version by Joachim Goedhart (40)). Timeplots of difference illustrate how these effects evolve over time, aiding in assessing their biological relevance within the context of the study. Such plots enable experimentalists to infer insight by evaluating the effect size. Finally, existing tools do not take into account cellular plasticity in the form of behavior switching as a rich differentiator of cell behaviour (38, 39). Given that each cell in each cluster is defined using 29 individual metrics and no user thresholding, a cluster switch based on this definition contains within it more information than one that is based on thresholding a single metric, thus providing a more reliable way to measure complexity or plasticity of behaviour compared with current methods in the field. Therefore, cellPLATO offers a new way to define clusters of behaviour, to understand the nature of cells in each cluster intuitively, and to understand the switching between clusters of any population of cells. This tool enabled us to analyze imaging data from 10-hour timelapses of enriched human NK cells from peripheral blood interacting with integrin ligands ICAM-1 or VCAM-1 in the presence of IL-15. We found eight distinct single timepoint behaviours, which were combined into 4 different behavioural trajectories over time. Each of these trajectories had unique responsiveness to IL-15 and differed in their frequencies between ICAM-1 and VCAM-1.

### Patterns of cell shape and motility

Initially, we used cellPLATO’s measurement and conditional comparison modules to plot population-level data. We found that NK cells on ICAM-1 and VCAM1 react strongly and acutely to the addition of IL-15 by increasing their migration speed and cell area. Interestingly, NK cells on ICAM-1 underwent these increases more rapidly than those on VCAM-1. NK cells on ICAM-1 also had a much greater area with similar cell migration speed compared with those on VCAM-1, indicating that though NK cells increase their migration speed on both surfaces, they might be doing so using differing forms of migration. We subsequently used cellPLATO to investigate sub-populations of cells responding differently to IL-15 on each surface.

Using the tool’s dimensionality reduction, trajectory designation, fingerprinting, and single cell de-abstractification features, we identified 8 single timepoint behaviours (clusters) defined by their combinatorial morphology and migration characteristics. The proportion of cells in each cluster differed between ICAM-1 and VCAM-1. Specifically, ICAM-1 induced the prevalence of cells with larger areas, medium polarization, and directed migration of medium speed. Conversely, VCAM-1 induced the prevalence of very fast cells with high polarization and less directed migration. These differences between modes of migration are consistent with ICAM-1-induced crawling behaviour in other lymphocytes (12, 47) and VCAM-1-induced migration in which less of the cell body is in direct contact with the substrate (48, 49). Both modes of migration can be visualized on developmentally supportive stromal cells that express a mix of ICAM-1 and VCAM-1 (2) and stromal cell-derived extracellular matrix containing collagen and fibronectin (5).

While information about cell dynamics was extracted using clustering of single timepoint behaviors, we further sought to understand how cells could move between clusters throughout the time of imaging. Using trajectory analysis, we uncovered 4 trajectories of cell behavior represented by different combinations of the single timepoint behaviours that were present in different proportions on ICAM-1 or VCAM-1. Cells adopted 1) stationary, semi-quiescent behaviour, 2) highly plastic, exploratory migratory behaviour, 3) delayed activation and then consistent migration behaviour or 4) oscillatory ‘stop and go’ confined migration type behaviour. Each surface had a similar proportion of cells with delayed activation that initiated migration after 80 minutes, while on VCAM-1 a subset of very exploratory cells travelling far distances and undergoing frequent shape changes was increased compared with ICAM-1. Oscillatory ‘stop and go’ confined (trajectory 3) subtype was much increased in the case of ICAM-1. The ICAM1 condition had fewer stationary, quiescent cells that didn’t activate than VCAM-1. The most plastic subsets of cells were increased on VCAM-1 compared with ICAM-1. The morphology of these cells changed rapidly and frequently between all 8 single timepoint clusters as the cells migrate (Movies 3-6). In contrast, intermediate stop and go type migration was more prevalent in the NK cells on ICAM-1.

Delayed activation of cells happened on both surfaces in a subset of cells (1∽8% on ICAM-1 and 25% on VCAM-1). Upon beginning their migration, such cells then underwent confined exploration, or move to a nearby area and searched their environment from a tethered position by short movements and polarization/re-orientation. The presence of this subset of cells in conjunction with the presence of highly exploratory cells, as well as a subset that do not respond to IL-15 and remain stationary (but alive), indicates that some cells are pre-tuned to respond quickly to IL-15, whereas others need to upregulate components that are required for migration. The reaction to IL-15 of such cells being delayed may indicate a recruitment of stored integrin ligands to the cell surface or the genetic upregulation of such ligands or other migration components. cellPLATO allowed us to detect these two populations in a data-driven, less biased manner than manual curation, allowing us to separate cell subpopulations from heterogeneous populations based on phenomics and morpho-kinetic features.

Finally, by imaging cells in the presence and absence of IL-15, we confirmed the phenomena described above in a third donor. We also showed that IL-15 reduced the population of quiescent cells and increased the other more migratory and exploratory populations. We saw acute increases in behavioral plasticity after the addition of IL-15 which were not present in cells that did not receive IL-15. We also describe subsets that respond quickly or slowly to IL-15 addition, confirming the acute effects of IL-15 on spontaneous migration in NK cells. We cannot rule out that cell division, known to be upregulated by IL-15 and not quantified here, generates additional cells tuned for one type of migration or another on either ICAM-1 or VCAM-1. Together, these data demonstrate that both integrin ligand and IL-15 signaling can modulate heterogeneous modes of migration of mature NK cells.

### Limitations and next steps

Trajectory analysis allowed us to formally identify 4 subsets of behaviour in our enriched, yet heterogeneous, NK cell cultures, which we have described informally previously on stromal cell monolayer systems using manual and semi-automated tracking approaches (5, 50). The hardness of the glass surface (51–53), the lack of integrin crosstalk produced by multiple ligands (54–56), the lack of shear flow (57), and the 2D nature of the system used here (53) limited our biological interpretations to those specific to these surfaces. However, here we provide new information about the acute response of subsets of mature NK cells to IL-15 on ICAM-1 and VCAM-1 on a stiff glass surface. Future studies could focus on linking these cell migration phenotypes with previously identified phenotypic subsets of mature NK cells. Our previous studies have identified differential modes of migration between CD56^bright^ and CD56^dim^ NK cell subsets (50), and these subsets also have differing expression of β1 and β2 integrins that mediate binding to ICAM-1 and VCAM-1 (9). Understanding how integrins, cytokine activation, and developmental subsets interact to regulate motile scanning of NK cells is an ongoing area of interest. These analyses were driven by the development of cellPLATO. Its adaptable and modular format provides a good basis for community-driven development and usage, as it is amenable to the addition of new metrics that do not affect the downstream flow of analysis and can be implemented simply. In the future, cellPLATO will be extended to perform shape and motility analysis on data acquired at different magnifications and timescales and incorporate 3D measurements, network analysis, fluorescent reporters of cell state (58, 59) and cell lineages or cell divisions (31–33, 44, 45). Access to higher resolution cellular features in small subsets of cells placed within a population will be a powerful way to link molecular behaviours to populations of cells. Together, cellPLATO provides a user-friendly platform to analyze heterogeneity in complex cellular populations, is open source and adaptable to different inputs, and is under continued development. The current version of cellPLATO is available for download with complete instructions for usage on Github with interactive Jupyter Notebooks provided.

## Methods

### Primary human NK cell culture

Blood samples were collected in sodium heparin tubes (BD Vacutainer) from healthy donors at Columbia University Irving Medical Center. All samples were obtained with informed consent under guidelines established by the Declaration of Helsinki. Primary NK cells were isolated from peripheral blood by negative selection using RosetteSep human NK enrichment cocktail (Stemcell Technologies, 15065). Blood was layered onto a Ficoll-Paque (Fisher Scientific, 45-001-750) density gradient followed by centrifugation at 1200 x g for 20 minutes with no brake. NK cells were collected from the density gradient interface and were placed into PBS at RT, before being centrifuged again for 10 minutes at 300 x g with a low brake. The cell pellet was then resuspended in 20 ml of warm cell culture medium: R10 (RPMI 1640 (Thermo Fisher, 11875135), 10% heat-inactivated fetal bovine serum (FBS, Gemini, 900-108-500), 10 mM HEPES (Fisher Scientific, 15630130), 100 U/mL Penicillin-Streptomycin (Fisher Scientific, 15140163), 1 x MEM Non Essential Amino Acids (Thermo Fisher, 11140076), 1 x Sodium Pyruvate (Fisher Scientific, 11360070), 1 x L-Glutamine (Glutamax, Thermo Fisher, 35050079), prior to centrifugation at 120 x g for 10 minutes with a low brake. Supernatant was discarded and cells were resuspended in R10 to a concentration of 2.5 x 10^6^ cells/ml. Cell solution was transferred to a 48 well plate and placed into an incubator at 37 °C with 5% CO2 for 10 minutes prior to use in microscopy or flow cytometry experiments.

### Preparation of integrin ligand coated chamber slides for microscopy

40 μl of 0.01% poly-L-lysine in di-H2O was added to each port of a μ-Slide VI 0.5 Glass Bottom chamber slide (Ibidi, 80607) which was incubated for 1h at RT. Chambers were washed 5 times with 120 μl of PBS by addition to one port and removal at the opposite port. At the final wash, all liquid (160 μl) was removed before addition of integrin ligands. 160 μl of 1 μg/ml Fc-ICAM-1 (Recombinant Human ICAM-1/CD54 Fc Chimera Protein, CF, R&D, 720-IC-200) or 1 μg/ml Fc-VCAM-1 (Recombinant Human VCAM-1-Fc Chimera, Biolegend, 553706) was added to the chamber slide, which was then incubated at 37 °C with 5% CO2 for 2 hours. Migration media (R10 media with 1.2% FBS) was pre-equilibrated in the incubator at 37 °C with 5% CO2 for 2 hours prior to its use for washing the imaging plate by adding 120 μl of media through one port, removing 120 μl of solution at the opposite port, repeated twice.

### Isolated NK cell preparation for fluorescence microscopy

NK cells were labelled with SPY650-DNA (Cytoskeleton, CY-SC501) and Cellbrite Steady 550 with Cellbrite Enhancer reagent (Biotium, 30107-T) by first making a 10X solution by diluting the three reagents 1:100 in R10 medium. IL-15 (Peprotech, 200-15) was added to this solution at 10X (0.1 μg/ml) for IL-15 conditions and the dye/cytokine solution was added to cells to a final concentration of 1X (10 ng/ml). The solution was mixed, and cells were incubated for 5 minutes at 37°C with 5% CO2.

### Timelapse confocal microscopy

μ-Slide channels were imaged using a confocal spinning disk microscope (Zeiss/3i/Yokogawa CSU W-1) equipped with a 37°C incubator supplied with 5% CO2 (OKOlab). Prior to adding the sample to the microscope, the microscope, stage, and insert were equilibrated for 2 hours to enhance stability and reduce sample drift during imaging. The sample was subsequently equilibrated on the stage for 30 minutes. Tiled (2X2) and montaged time-lapse images of a single focal plane in multiple wells were obtained using a 20X 0.8 NA air objective with a 40s interval between time frames. Imaging was carried out using a 561 nm laser at 8% power with 120 ms integration time and a 640 nm laser at 9% power with 150 ms integration time. Imaging was performed for a total of 900 frames (600 min; 10 hours) using an sCMOS camera (Teledyne Photometrics Prime 95B). To control for potential toxicity of membrane and nuclear dyes and their exposure to laser light, brightfield imaging in cells free of dyes was carried out simultaneously to fluorescent imaging in a test experiment and no change in behaviour was observed.

### Image processing

2X2 tiled timelapse data was montaged in Slidebook 6.0 software (Intelligent Imaging Innovations) using a 5% image overlap before being exported as an image sequence of TIF files. The bioformats (60) plugin in Fiji (61) was used to open the sequence as a TIF stack. Images were cropped and then exported again using bioformats as an image sequence of a separate numbered TIF per timepoint. Each TIF contained 2 channels, where C1 was the membrane channel (SPY650-DNA), and C2 was the nuclear channel (Cellbrite Steady 550). We recommend using a file tree system as follows: all data is stored within a master folder, which contains a separate folder for each condition (ICAM-1 or VCAM-1 in this experiment). Inside each conditional folder, there is a separate folder for each replicate (i.e. donor 1, donor 2 and donor 3). Timelapse movies are saved as image sequences inside each replicate folder.

### Cell segmentation

The human in the loop feature of Cellpose 2.0 (27, 62) was used to retrain the existing neural network ‘Cyto2’. 10 example imaging frames were randomly selected from different conditions and replicates and were opened using the Cellpose GUI. A diameter of 28.6 was used with Cyto2, available in the model zoo, to successfully identify and segment most cells within the field of view. Each section of the field of view was examined and errors in segmentation were corrected manually by deleting the existing mask and repainting one or several manually. Errors often occurred tightly adjacent cells or in stretched out cells with both thick and thin regions. The corrected masks were used to retrain a new model called ‘Cyto2plus’ that was used for downstream analysis. The anaconda command line was used to batch segment all microscopy data within the master folder using PyTorch and an RTX 3090 GPU (NVIDIA).

### Cell tracking

Tracking of the segmented cells was performed using btrack (31). Hyperparameters and priors for the Bayesian model were optimized by changing each parameter manually and checking the fidelity of example tracks versus manually tracked ground truths. More detail about model optimization can be found in the btrack documentation. btrack was run using Jupyter notebook (included with the cellPLATO package) in batch on the segmented masks in the master folder. Segmentations, xy coordinates, and track IDs for each cell over time are saved into a separate h5 file per replicate.

### cellPLATO analysis

cellPLATO can be installed using pip and full instructions are available on Github. The software was built to be used by experimenters with no Python expertise and was tested and developed iteratively with users. Users input experimental parameters into a single config.py file including at a minimum 1) the path to the master folder containing all data, 2) the names of the conditions to be included in the experiment, 3) the pixel size and 4) the time resolution of the experiment. More advanced parameters include 1) the size of the sliding time windows used to normalize some of the distance dependent metrics (i.e. cumulative length – in this case it was set to 5 minutes), 2) the filters placed on the data to exclude small/fast objects such as debris from the analysis, and cells that were tracked for fewer frames than the minimum time window. Here only cells with an area of >50 μm^2^ and with at least 8 timepoints were included. This choice of time window was based on an average NK cell migration speed of 3∽ μm/min and cumulative length 1∽6 μm per time window reflecting the average length of two NK cells. Users should tune the size of the time window to their data. After completing and saving the config.py file, users can run the entire analysis using a Jupyter notebook. The notebook runs sequentially the measurement phase, the dimensionality reduction, clustering, and disambiguation phase, followed by trajectory analysis. Cluster membership scoring is calculated using a silhouette score ranging from -1 to 1: a high score indicates that a given data-point is well matched to its own cluster, and poorly matched to other clusters. The dataset used for this manuscript consisted of 3 separate blood donors. For each blood donor, the NK cells imaging on different integrin ligands in the presence or absence of IL-15 was performed simultaneously to make time-based comparisons between conditions. Each imaging timelapse contained 900 frames of size 2030X2030 pixels. Total time for full analysis was 7 hours using a mid-range CPU. Statistical testing was performed using Mann Whitney U for non-parametric data and ANOVA for parametric data to produce P-values, combined with calculation and plotting of the effect size distribution (also known as plots of differences41). Crossing of the effect size distribution with the control line is an indication of statistical inconsequence (46).

### Flow cytometry

NK cell enrichment was validated using a 5-color flow cytometry panel, confirming that >85% of enriched cells were CD56+CD3CD20CD14 NK cells (Supp. Fig. 1). Cells were incubated with antibodies for CD3 (clone SK7, BV421, Biolegend, 1:200), CD14 (clone M5E2, BV421, Biolegend, 1:200), CD19 (clone HIB19, BV421, Biolegend, 1:200), CD56 (clone HCD56, BV605, BioLegend, 1:100), CD18 (clone CBR LFA-1/2, AF647, Biolegend, 1:100), and CD29 (clone TS2/16, AF488, Biolegend, 1:100). A Zombie NIR Viability Dye was used for live/dead confirmation (1:200, Biolegend). Flow cytometry was performed on a Novocyte Panteon. All flow cytometry data analysis was performed with FlowJo (BD Biosciences) software.

## Code availability

The latest version of cellPLATO and supplementary movies described in this preprint can be found at github.com/Michael-shannon/cellPLATO. The version used for this paper can be found on Zenodo at doi:10.5281/zenodo.10056150.

## ACKNOWLEDGEMENTS

This work was supported by NIH-NIAID R01AI137073 to EMM. We also thank Paul Hsieh-Fu Tsai and Amy Shen at Okinawa Institute of Science and Technology for assistance in integrating Usiigaci with cellPLATO.We thank Ricardo Henriques for generating the BioRxiv template used for the preprint.

## Notes

https://github.com/Michael-shannon/cellPLATO

doi:10.5281/zenodo.10056150

### Competing Interest Statement

The authors have declared no competing interest.

### Summary of Updates

Title change and minor text edits.

