## Supplementary material for "cellPLATO: an unsupervised method for identifying cell behaviour in heterogeneous cell trajectory data": Table 1

| Cluster ID | Characteristic 1 | Characteristic 2 | Characteristic 3 | Morphology | Migration |
|---|---|---|---|---|---|
| **0** | Medium aspect ratio | Medium outreach ratio | Low MSD | Circular | Very low |
| **1** | Medium aspect ratio | High endpoint directionality ratio | High euclidean distance | Slightly less circular | Medium |
| **2** | Medium aspect ratio | High endpoint dir ratio | High euclidean distance | Slightly less circular | Medium |
| **3** | Medium aspect ratio | High endpoint dir ratio | Medium area | More spread, less circular | High |
| **4** | High aspect ratio | High Eccentricity | Medium area | Polarized | Medium |
| **5** | High aspect ratio | Medium outreach ratio | High euclidean distance | Polarized | Med/High |
| **6** | High aspect ratio | High endpoint directionality ratio | High euclidean distance | Polarized | Very straight and high |
| **7** | High aspect ratio | High euclidean distance | Low outreach ratio | Polarized | Very fast, irregular |

**Table 1: The top three characteristics of cells belonging to 8 distinct behavioural clusters described by morphological/motility measurements.** Peripheral blood derived IL15 treated cells NK cells migrating on a surface of ICAM-1 or VCAM-1**.** Rows of table coloured by cluster ID. N = 14895 cells from 1 blood donor.
