## Supplementary figures for "cellPLATO: an unsupervised method for identifying cell behaviour in heterogeneous cell trajectory data"

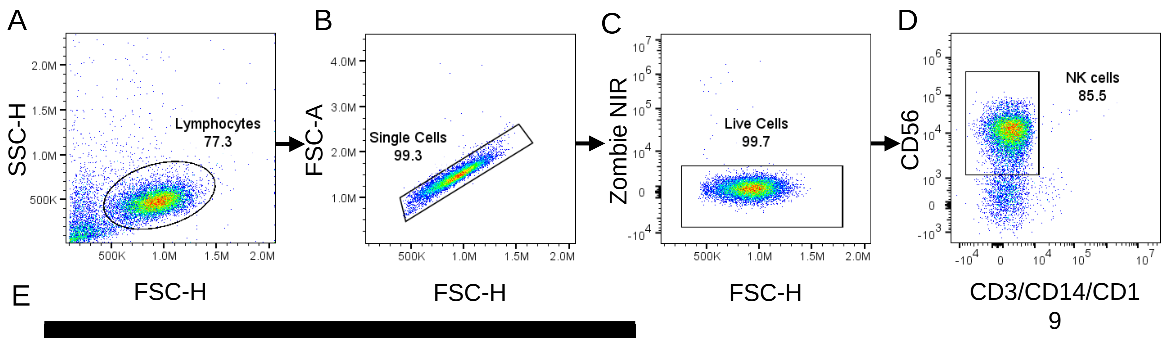


E

| Fluorophore | Marker | Clone; Source; Dilution |
|---|---|---|
| BV421 | CD3 | SK7; Biolegend; 1:200 |
| BV421 | CD14 | M5E2; Biolegend; 1:200 |
| BV421 | CD19 | HIB19; Biolegend; 1:200 |
| BV605 | CD56 | HCD56; Biolegend; 1:100 |

**Supplementary figure 1. Flow cytometry of enriched NK cells.** NK cells were enriched from human peripheral blood as described in Methods. Flow cytometry analysis was used to confirm enrichment for NK cells. A) The lymphocyte population was defined by low granularity (SSC-H) and high forward scatter by height (FSC-H). B) Within this population, single cells were defined by height (FSC-H) being lower than area of each detection peak (FSC-A). C) Zombie NIR was used to gate on living cells within the single cell population. D) CD56 positive and CD3/CD14/CD19 negative cells were defined as NK cells. E) Table of fluorophores and antibody clones used for flow cytometry. Representative plots from one donor are shown.

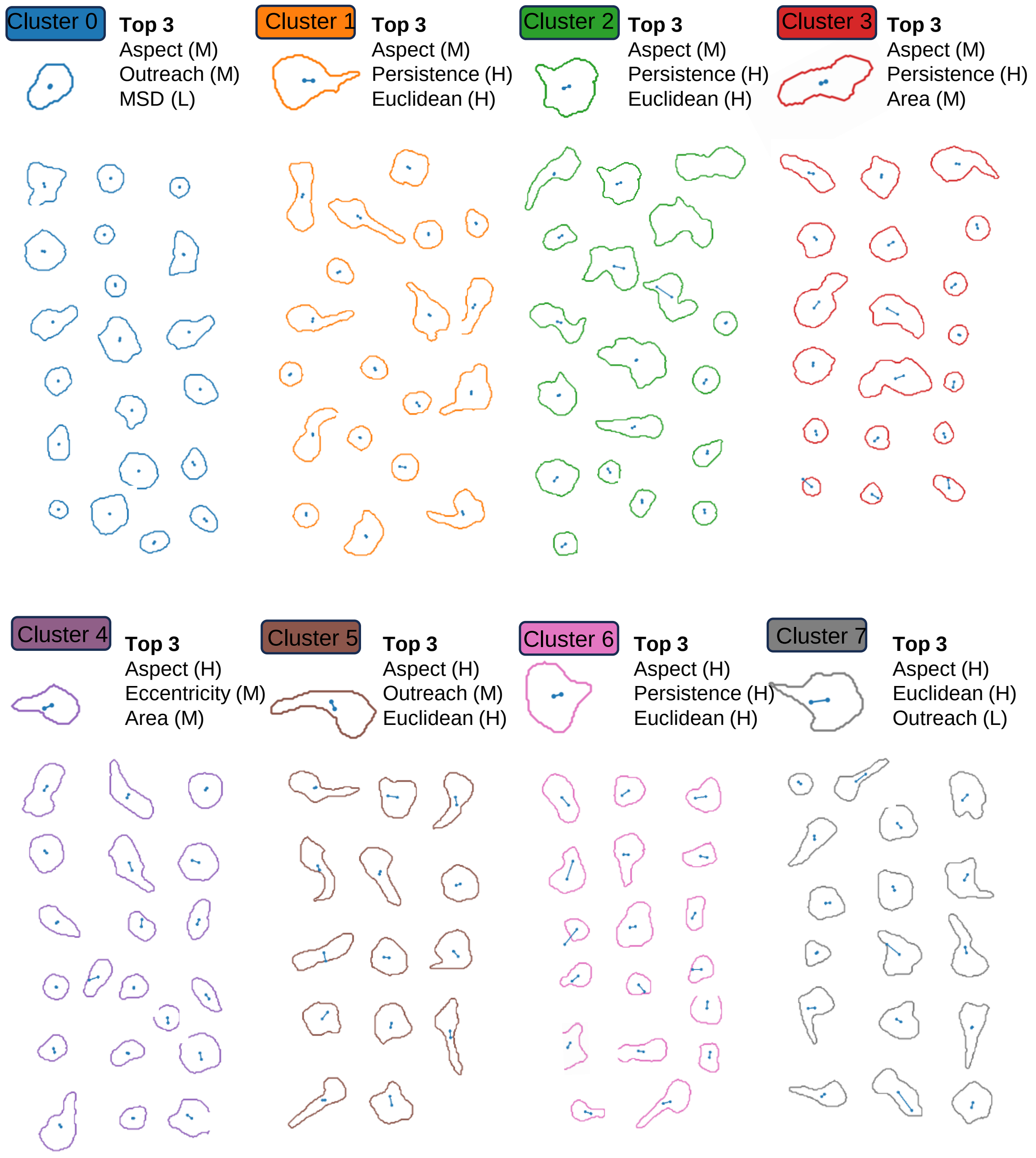


**Supplementary figure 2: Gallery of exemplar cells from each single timepoint behavioural cluster.** A) Exemplar cell contour and tracks from cluster 0; top 3 contributory metrics labelled high, medium, or low (H, M, L). B to H) Clusters 1 to 7. Exemplars taken from n = 14,985 cell tracks, 1,036,092 individual datapoints.

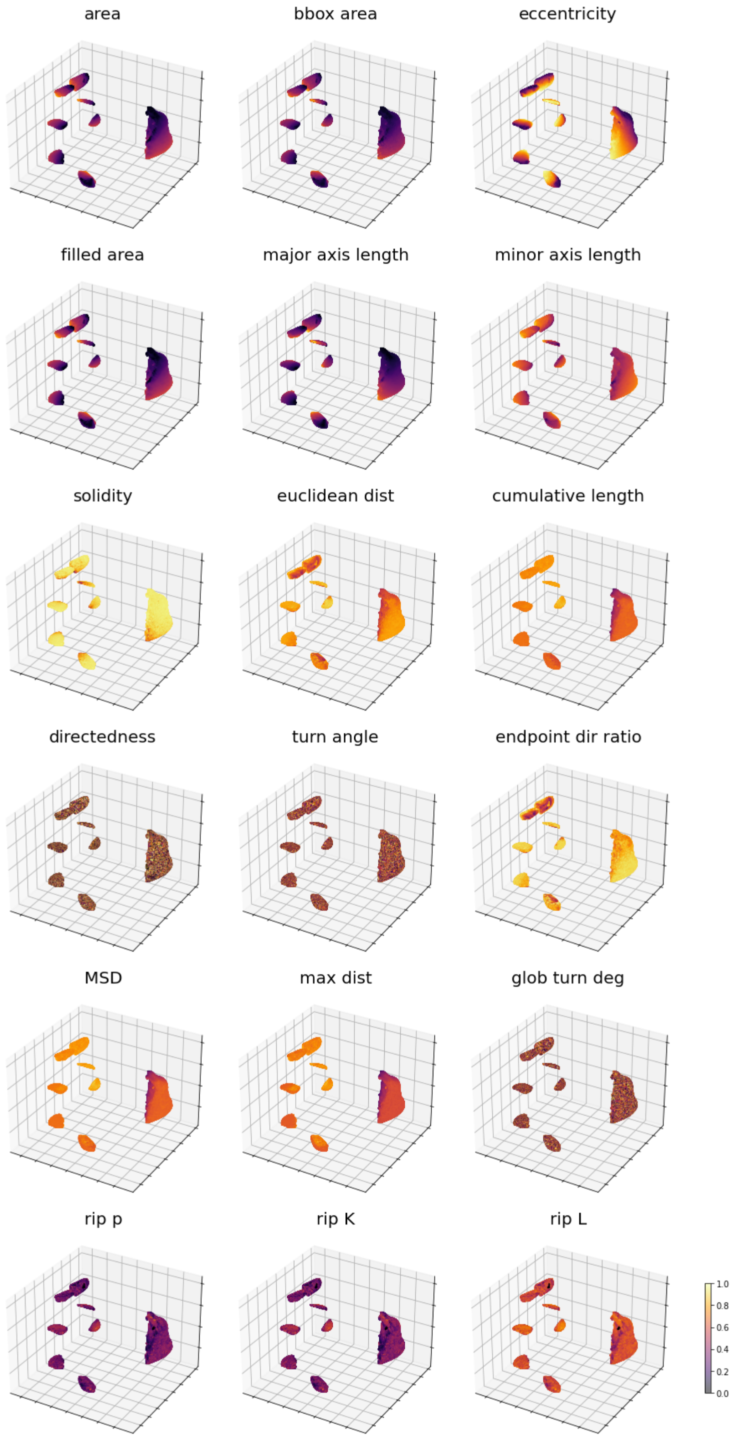


**Supplementary figure 3. UMAP plots coloured by contribution of each factor.** Plots of each metric with scaling used for dimensionality reduction (log2 and minmax). Each datapoint coloured by magnitude of each factor. n=14,825 cells from 1 donor.

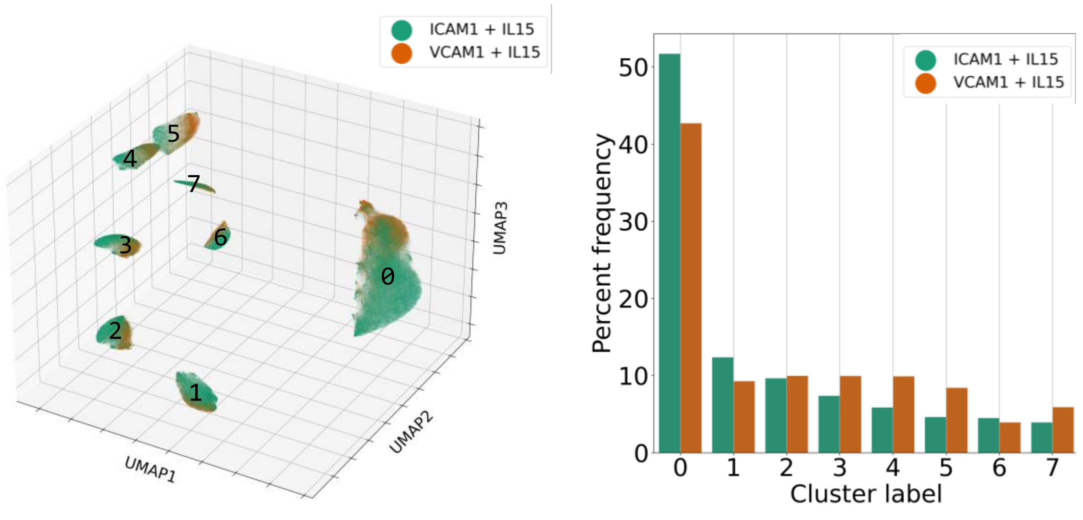


**Supplementary figure 4. NK cells occupy clusters differentially depending on their response to ICAM-1 or VCAM-1.** A) 3D UMAP coloured by condition (ICAM-1, green; VCAM-1, orange). B) Percent occupancy of cells from each condition in each cluster. Peripheral blood derived IL-15 treated cells NK cells migrating on a surface of ICAM-1 or VCAM-1. n = 14,825 cells from 1 blood donor.

| ClusterID | Metric | Median | Category |
|---|---|---|---|
| 0 | Aspect | 1.1817 | Medium |
| 0 | Outreach Ratio | 0.2508 | Medium |
| 0 | MSD | 0.2553 | Low |
| 1 | Aspect | 1.3179 | Medium |
| 1 | Endpoint Dir Ratio | 0.3164 | High |
| 1 | Euclidean Dist | 2.663 | High |
| 2 | Aspect | 1.3422 | Medium |
| 2 | Endpoint Dir Ratio | 0.3017 | High |
| 2 | Euclidean Dist | 3.3373 | High |
| 3 | Aspect | 1.3731 | Medium |
| 3 | Endpoint Dir Ratio | 0.3077 | High |
| 3 | Area | 136.3985 | Medium |
| 4 | Aspect | 1.3944 | High |
| 4 | Eccentricity | 0.6969 | Medium |
| 4 | Area | 130.3428 | Medium |
| 5 | Aspect | 1.4072 | High |
| 5 | Outreach Ratio | 0.2372 | Medium |
| 5 | Euclidean Dist | 6.827 | High |
| 6 | Aspect | 1.3929 | High |
| 6 | Endpoint Dir Ratio | 0.5095 | High |
| 6 | Euclidean Dist | 14.8644 | High |
| 7 | Aspect | 1.4194 | High |
| 7 | Euclidean Dist | 9.674 | High |
| 7 | Outreach Ratio | 0.2206 | Low |

**Supplementary table 1: Average characteristics for each of top three characteristics of cells belonging to 8 distinct behavioural clusters described by morphological and motility measurements.** Peripheral blood derived IL-15 treated cells NK cells migrating on a surface of ICAM-1 or VCAM-1**.** n = 14,825 cells from 1 blood donor.

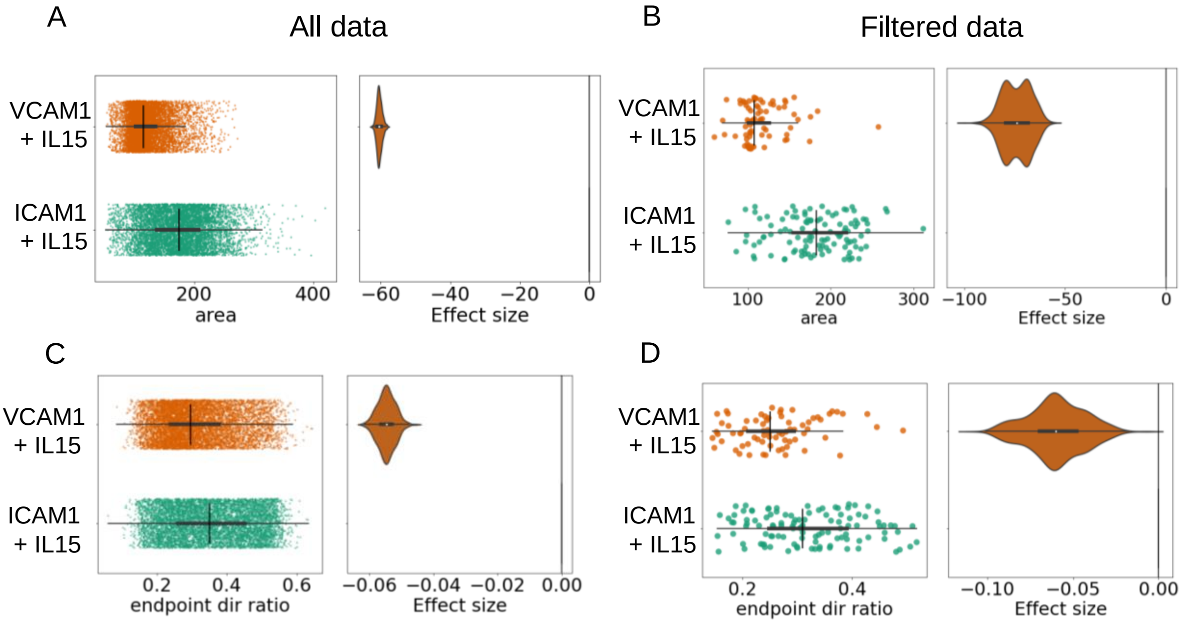


**Supplementary figure 5. Cell area and endpoint directionality ratio trends are maintained in filtered data compared with full data.** Data was filtered to only include cell tracks between 200 to 220 timepoints in length to allows for sequence similarity measurement. A) Area of cells (plots of difference) for full data and B) filtered data. C) Endpoint directionality ratio (persistence length) for full data and D) filtered data. Green – NK cells on ICAM1; Orange – NK cells on VCAM1. N= 14825 cells in full data, 187 cells in filtered.

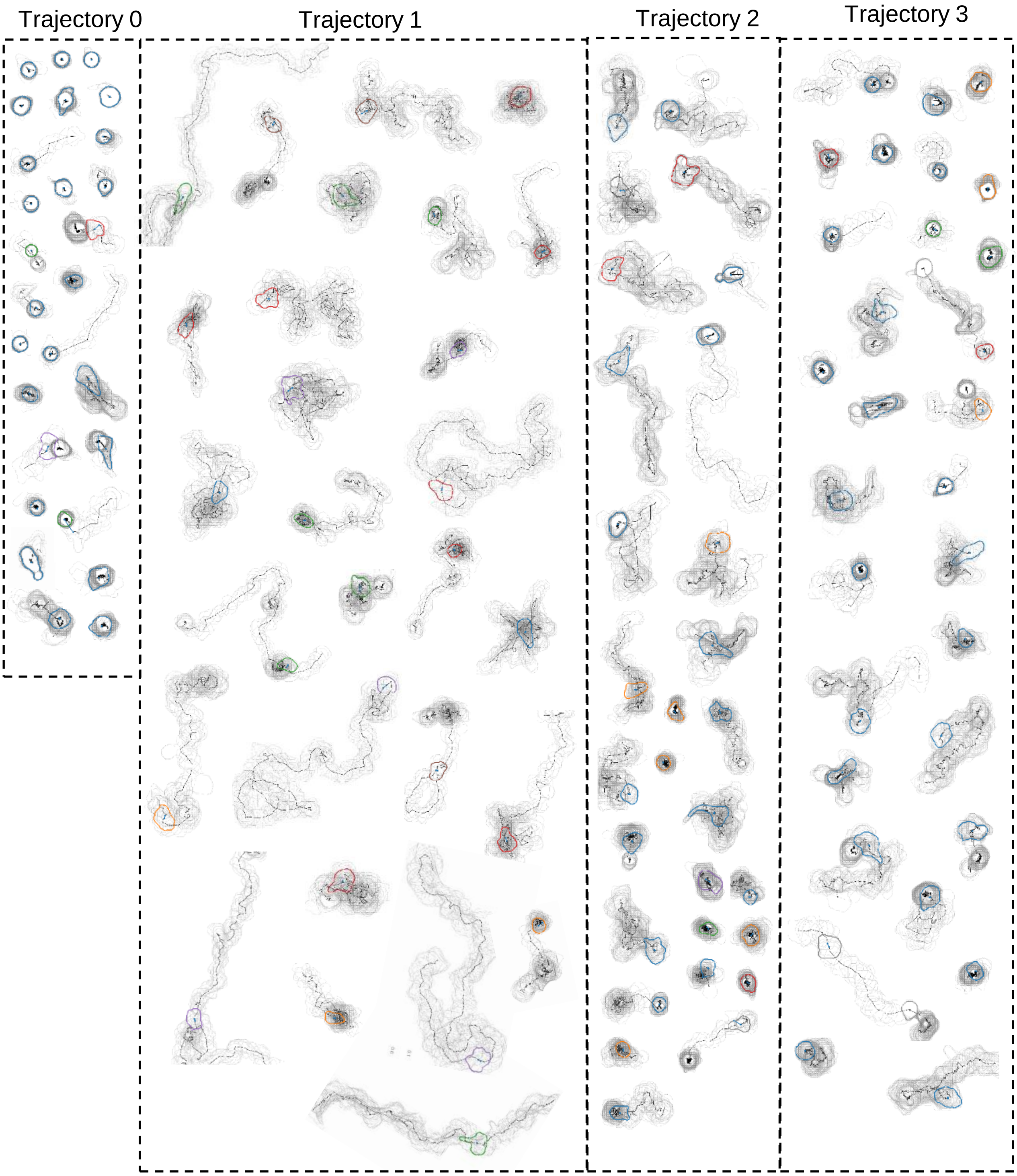


**Supplementary figure 6. Gallery of behavioural trajectory IDs as contour plots and time traces.** Representative cells assigned to each trajectory ID. Representative cells taken from n=183 cells from donor 1.

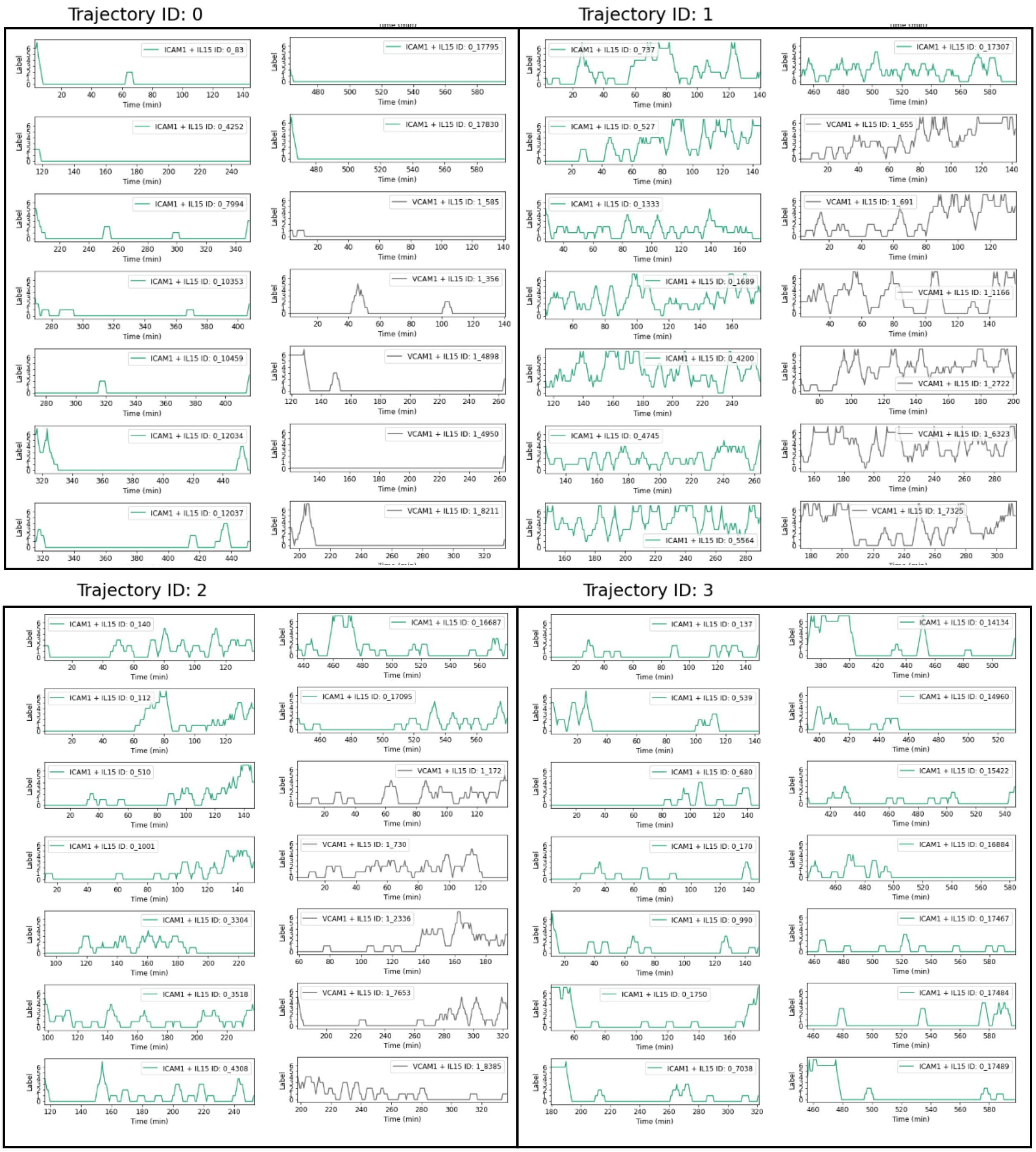


**Supplementary figure 7. Behavioural trajectory IDs as plots of behavioural cluster ID over time.** Additional representative examples of cluster ID (label) over time for each trajectory ID (green, ICAM-1; grey, VCAM-1). Representative cells from n = 183 cells from donor 1.

**Supplementary movies are available here:**

https://drive.google.com/drive/folders/1wvCbWoywRdk0OWhcwwJhiIdAEz4yTKHt

**Supplementary Movie 1: Timelapse micrographs of purified human NK cells migrating on ICAM1 or VCAM1 in the presence of IL15.** Top panels show stitched fields of view of NK cells from donor 1 migrating on ICAM1 (left) or VCAM1 (right). Bottom panels show zoomed regions denoted by black boxes for ICAM1 (left) and VCAM1 (right). Images are pseudocolored with yellow denoting cell cytoplasm labelled with Cellbrite Steady 550 and blue denoting nuclei labelled with SPY650-DNA. Micrographs are representative of several fields of view of cells from 3 donors imaged on each surface for 10 hours.

**Supplementary Movie 2: Segmentation and tracking of purified human NK cells migrating on ICAM1 or VCAM1 in the presence of IL15.** Top panels show segmentation masks and tracks in stitched fields of view of NK cells from donor 1 migrating on ICAM1 (left) or VCAM1 (right). Bottom panels show zoomed regions denoted by black boxes for ICAM1 (left) and VCAM1 (right). Segmented masks are arbitrarily colored per ID per frame, and tracks are colored by cell ID. Masks and tracks are representative of several fields of view from 3 donors imaged on each surface for 10 hours.

**Supplementary Movie 3: Trajectory cluster 0 animation of contour and track matched with raw image data.** Several representative cells from trajectory cluster 0 extracted and displayed. For each cell, the left pane denotes their contours and tracks, where the colour of the contour denotes the single timepoint cluster ID and the black line represents the track. To provide context, the right pane shows the cell in the raw imaging data to which each contour/track relates.

**Supplementary Movie 4: Trajectory cluster 1 animation of contour and track matched with raw image data.** Several representative cells from trajectory cluster 1 extracted and displayed. For each cell, the left pane denotes their contours and tracks, where the colour of the contour denotes the single timepoint cluster ID and the black line represents the track. To provide context, the right pane shows the cell in the raw imaging data to which each contour/track relates.

**Supplementary Movie 5: Trajectory cluster 2 animation of contour and track matched with raw image data.** Several representative cells from trajectory cluster 2 extracted and displayed. For each cell, the left pane denotes their contours and tracks, where the colour of the contour denotes the single timepoint cluster ID and the black line represents the track. To provide context, the right pane shows the cell in the raw imaging data to which each contour/track relates.

**Supplementary Movie 6: Trajectory cluster 3 animation of contour and track matched with raw image data.** Several representative cells from trajectory cluster 3 extracted and displayed. For each cell, the left pane denotes their contours and tracks, where the colour of the contour denotes the single timepoint cluster ID and the black line represents the track. To provide context, the right pane shows the cell in the raw imaging data to which each contour/track relates.
